# Building the vector in? Construction practices contribute to the invasion and persistence of *Anopheles stephensi* in Jigjiga, Ethiopia

**DOI:** 10.1101/2023.05.23.541906

**Authors:** Solomon Yared, Araya Gebresilassie, Esayas Aklilu, Elyas Abdulahi, Oscar D. Kirstein, Gabriela Gonzalez-Olvera, Azael Che-Mendoza, Wilbert Bibiano-Marin, Elizabeth Waymire, Jo Lines, Audrey Lenhart, Uriel Kitron, Tamar Carter, Pablo Manrique-Saide, Gonzalo M. Vazquez-Prokopec

## Abstract

*Anopheles stephensi* is a major vector of malaria in Asia and the Arabian Peninsula, and its recent invasion into Africa poses a significant threat to malaria control and elimination efforts on the continent. The mosquito is well-adapted to urban environments, and its presence in Africa could potentially lead to an increase in malaria transmission in cities. Most of the knowledge about *An. stephensi* ecology in Africa has been generated from studies conducted during the rainy season, when vectors are most abundant. Here, we provide evidence from the peak of the dry season in the city of Jigjiga, Ethiopia, and report the finding of *An. stephensi* immature stages infesting predominantly water reservoirs made to support construction operations (in construction sites or associated with brick manufacturing businesses). Political and economic changes in Ethiopia (and particularly the Somali Region) have fueled an unprecedented construction boom since 2018 that, in our opinion, has been instrumental in the establishment, persistence and propagation of *An. stephensi* via the year-round availability of perennial larval habitats associated with construction. We argue that larval source management during the dry season may provide a unique opportunity for focused control of *An. stephensi* in Jigjiga and similar areas.

## Introduction

Remarkable success in reducing malaria burden has been achieved in most African countries since the year 2000, thanks to the scaling-up of vector control tools (insecticide-treated nets and indoor residual spraying) and effective preventive and treatment drugs^1^. Increasing evidence suggests that rapid urbanization of Africa’s human population (driven primarily by rural-urban immigration) is also contributing to a reduction in malaria burden ^2-5^. Lower habitat suitability for *Anopheles spp*. breeding and improvements in housing within African cities reduce human-mosquito contacts and can lead to lower *Plasmodium* spp. inoculation rates compared to rural settings ^2-5^. Environmental management in the form of housing improvement has gained research interest due to its sustained effect on *Anopheles* spp. mosquitoes and its positive impact on livelihoods ^6,7^. The WHO calls this approach “building the vector out”, and involves the adoption of practices that range from improved housing structures to retrofitting eave tubes and other approaches to limit mosquito entry indoors ^8^. This approach is also seen as a novel aspect of malaria control in urban settings, given most human population growth over the next century will be accounted for by the growing number of city dwellers ^9^.

As most sub-Saharan countries continue their push towards malaria elimination, a new threat has the potential to negatively impact decades of public health gains: the invasion and establishment of *Anopheles stephensi*, a malaria vector native to Asia, commonly found in cities throughout India, Iran, Pakistan, and the Arabian Peninsula ^10,11^. Since it was first detected in Africa in Djibouti in 2012^12^, *An. stephensi* has spread to Ethiopia^13^, Somalia^14^, Sudan^15^, Kenya, Nigeria and Ghana^16^. Niche modeling predicts suitable environmental conditions for *An. stephensi* establishment throughout tropical African cities, putting an additional 126 million people potentially at risk of malaria ^17^. Given the mosquito’s urban dependency and container larval breeding habits ^18,19^, rainfall alone was found to be a poor predictor of *An. stephensi*-driven malaria transmission ^20^. In Djibouti, a 2000x exponential increase in the number of malaria cases has been observed since the detection of *An. stephensi* ^11,16^. The contribution of *An. stephensi* to increases in malaria transmission outside of Djibouti has begun to be investigated, especially in light of the recent malaria outbreak. From 2018 to 2020 in Ethiopia, *Plasmodium vivax* was detected in wild-caught *An. stephensi* from the cities of Dire Dawa and Kebridehar (with infection rates of 0 ·5% and 0 ·3%, respectively)^19^ and *P. vivax* and *P. falciparum* infection recently reported in *An. stephensi* from Awash (2·8% and 1·4%)^21^. Furthermore, experimental membrane feeding experiments showed that field-caught *An. stephensi* from Ethiopia became significantly more infectious with local *P. vivax* and *P. falciparum* than *Anopheles arabiensis* (the primary malaria vector in Ethiopia), indicating that it is a highly competent vector for African *Plasmodium*^21^.

Given the entomological and epidemiological evidence gathered so far, the World Health Organization (WHO) launched a new initiative to stop the further spread of *An. stephensi* in the region that is based on a 5-pronged approach: 1. increasing collaboration; 2. strengthening surveillance; 3. improving information exchange; 4. developing guidance; and 5. prioritizing research^16^. To execute an effective plan for *An. stephensi* elimination, key sources of information about its biology and bionomics in its new habitats are needed, and vector control tools that are better suited for urban settings will need to be investigated.

Several studies (most of them from Ethiopia, cross-sectional and conducted during the rainy season) have characterized *An. stephensi* habitats and bionomics, with many knowledge gaps still remaining (e.g., ^18,19,21,22^). The evidence gathered so far shows that in the rainy season, *An. stephensi* larvae are found in a wide array of small and large artificial containers, ranging from large water cisterns to car tires and buckets ^18,19,21^. In addition to *Plasmodium* infection, such studies have characterized up to 48% human biting (14/29 mosquitoes) in Awash^21^ but low human biting (<1% human biting) in Dire Dawa and Kebridehar (where also a high frequency of domestic animal feeding was observed)^19^. Such discrepancies may have originated, in part, due to the opportunistic collection of adult mosquitoes in or near animal shelters. Indeed, the finding of *P. vivax* and *P. falciparum* infected mosquitoes can only be explained by human biting. Furthermore, given its egg-laying behavior (eggs that resist desiccation ^23^ and are laid in small containers), the fact that it bites humans not only at night when they are sleeping, and that it is found in urban and peri-urban areas, *An. stephensi* has more similarities with *Aedes aegypti* mosquitoes (which vector viruses such as dengue, chikungunya and Zika) than with other Anophelines ^24^. One of the many factors that remains to be studied is how is it that *An. stephensi* persists in Ethiopia and other African countries that have a prolonged dry season, as this period may offer unique opportunities for surveillance and control.

Here, we report novel findings on the habitat use of *An. stephensi* during the dry season in eastern Ethiopia. While increased focus on characterizing larval habitats in rainy periods can provide information of niche breadth for the species, our goal of focusing on the dry season was to explore possible windows for control in periods where the population size may be smallest.

### Dry season *An. stephensi* collections in Jigjiga, Ethiopia

From March 6-14, 2023, mosquito surveys were conducted in Jigjiga city (capital of Somali Region, Ethiopia, population ∼800,000) during the dry season. *Anopheles stephensi* was first detected in Jigjiga in 2018^18^, and has persisted in the city since then despite a harsh dry season (the rainless period of the year lasts for ∼3 months). Molecular analysis of cytochrome oxidase subunit I (*COI*) and cytochrome B gene (CytB) shows Jigjiga as one of the locations with highest diversity, suggesting it was likely an early introduction point of *An. stephensi* into Ethiopia^25^. Jigjiga is of relevance because of its large population size, rapid urbanization, and connection to other malaria-endemic regions and the port of Berbera in Somaliland.

We employed methods developed for standard larval and pupal sampling of container breeding mosquitoes^8^, that included collecting all the larvae and pupae in small water holding containers and using dippers and large fish nets to sample large water-holding habitats. All the larvae and pupae were reared to Adult at Jigjiga University Entomology Laboratory. The emerged *Anopheles* spp. adults were identified to species level using standard keys as well as molecular means. From a total of 60 potential larval sites with water that were sampled across the city, we identified a major habitat consistently positive for *An. stephensi* larvae and pupae during the dry season: man-made pits related with construction operations (Fig 1A). We term such habitats ‘construction pits’, as they were primarily built for the storage of water in construction sites or in small-scale brick manufacturing businesses (Fig 1B). *Anopheles stephensi* positivity in construction pits was 62·5%, compared to 5 ·9% in water cisterns made of cement and 0% in 200L plastic drums (Fig 1B). All abandoned tires sampled did not contain any water. Interestingly, from all the sites that we found positive for *An. stephensi* larvae and pupae, 63.6% of them also had *Culiseta* spp. and *Aedes* spp. larvae in them, whereas only 18·2% and 9·1% sites positive for *An. stephensi* were co-habited by *Culiseta* spp. or *Aedes* spp. only, respectively (Fig 1C).

**Figure 1.**
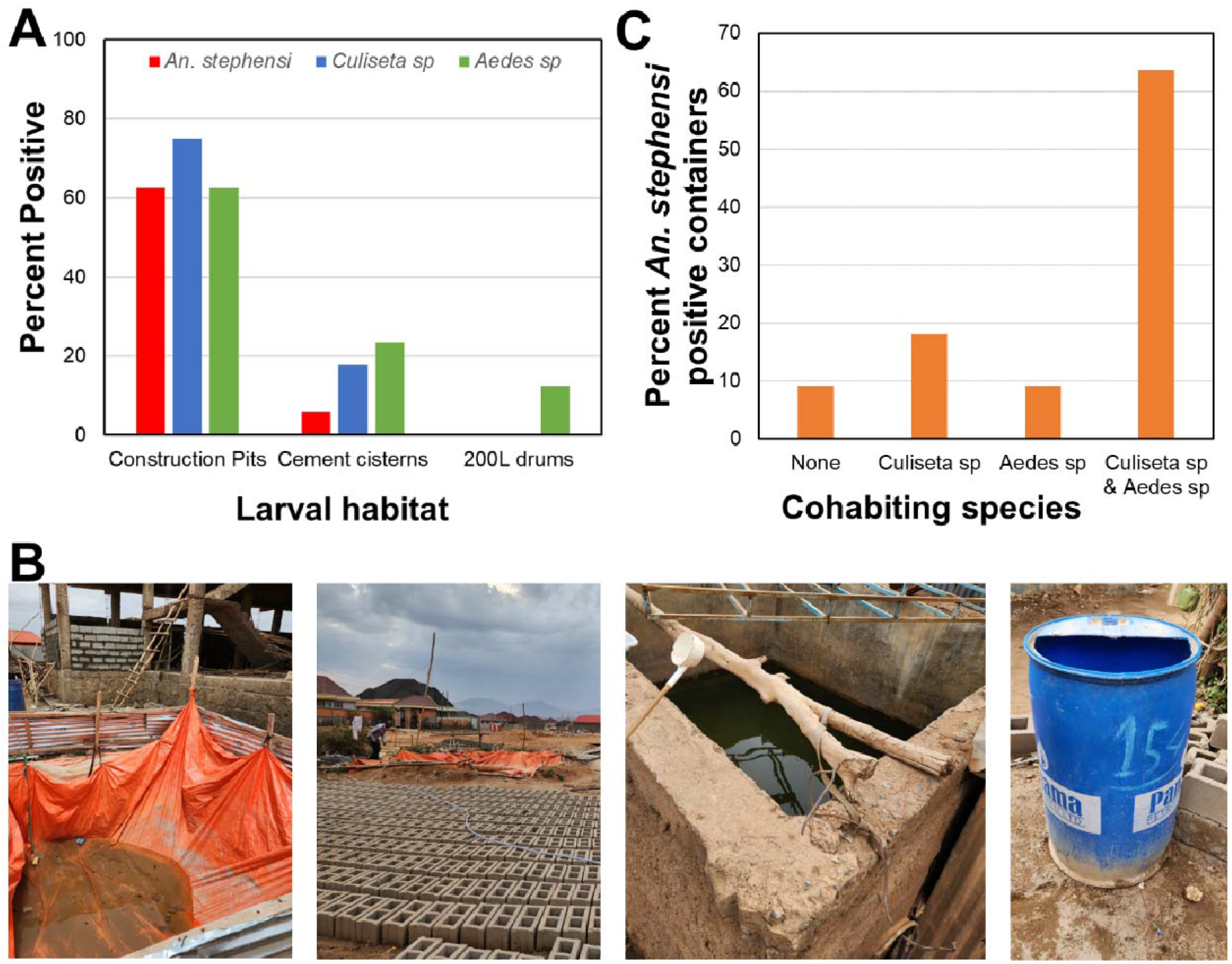
(A) Positivity of *Anopheles stephensi* immature stages in the only containers found with water during the dry season survey of 2023 in Jigjiga, Ethiopia, and stratified by species or genus of mosquito found. (B) Examples of sampled habitats, construction pits associated with house construction (left), a construction pit associated with brick manufacturing (center left), a cement cistern (center right) and a 200L plastic drum (right). (C) Species cohabiting with *An. stephensi* in positive containers (*Culiseta* represents *C. longiareolata*, whereas *Aedes* represents *Ae. hirsutus*).

A subset of 20 adult emerging from the pupae collected in construction pits and visually identified with standard keys was molecularly confirmed to be *An. stephensi* using an allele specific PCR and the sequencing of *ITS2* and *COI* loci^13^. While *ITS2* haplotypes were all identical for the *An. stephensi* samples, three *COI* haplotypes were detected: Hap 1 (7/14), Hap 2 (6/14), and Hap 3 (1/14) (using Carter et al 2021^25^ haplotypes designations), mostly consistent with previous studies. Notably, the presence the *COI* Hap 1 (common to South Asia and detected in northern Ethiopia and Djibouti) supports the notion of Jigjiga’s connectivity with regions outside of the continent with long-established *An. stephensi* populations and as a likely entry point for *An. stephensi* into the southern part of the country.

After the molecular confirmation of *An. stephensi*, we used the GPS coordinates from the construction pits in Google Earth to identify them remotely, given their unique spectral signature (size, color contrast and presence of water). Interestingly, using a high-resolution satellite image taken in November 2022 (4 months prior to sampling) we not only were able to identify the positive construction pits (Fig 2A-B) but also extended our work to identify a total of 101 pits within a rural to urban swath of Jigjiga centered on the road connecting the city with Somaliland (Fig 2C). The density of pits per hectare within the swath did not follow a rural-urban gradient but concentrated in the center of the swath, an area of Jigjiga currently experiencing rapid construction and development (Fig 2D). Given the high positivity rate of construction pits we quantified (Fig 1), the density map in Fig 4 may be a good proxy for the distribution of *An. stephensi* distribution within the rural-urban swath.

**Figure 2.**
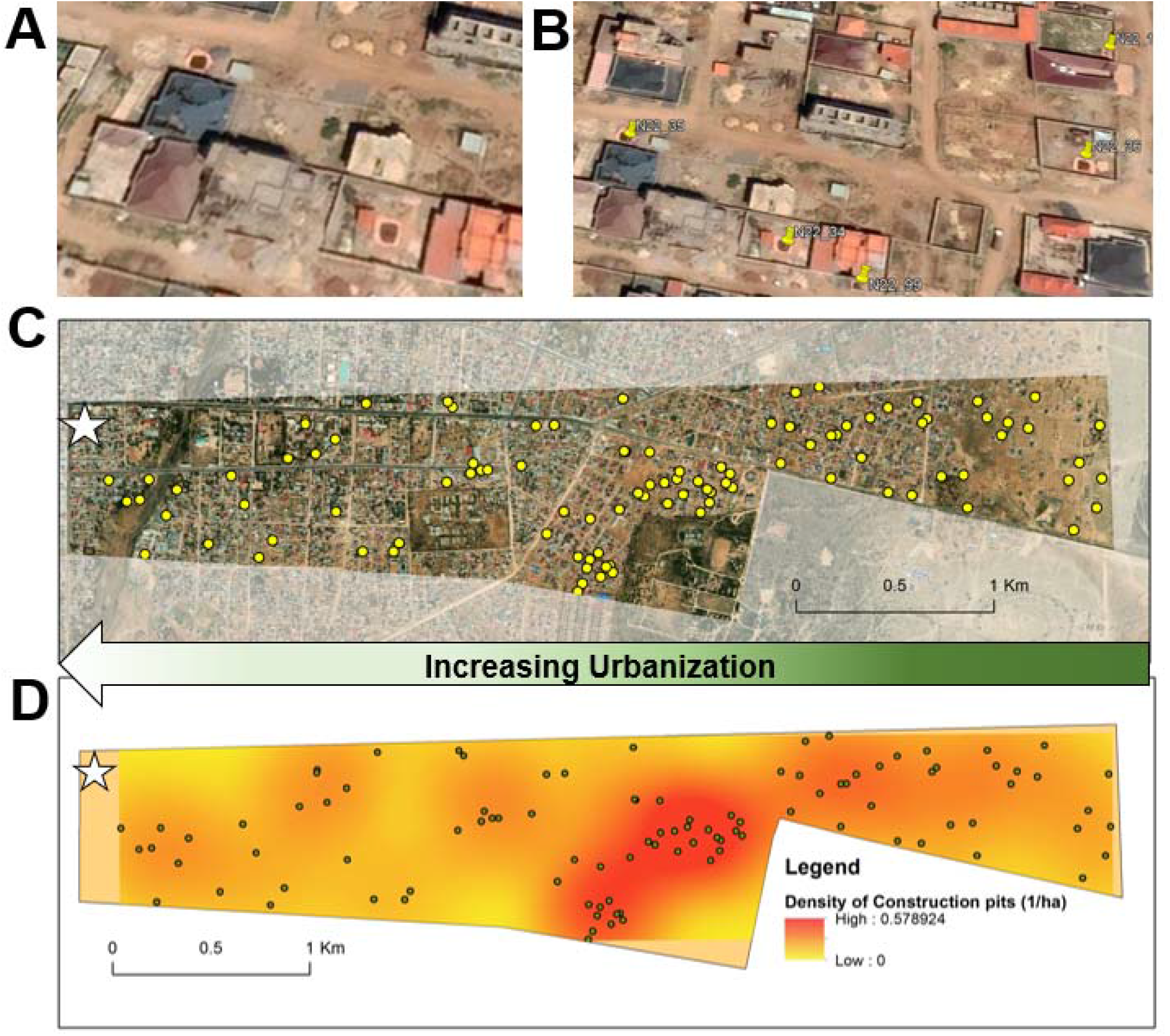
(A) Construction pits identified as positive for *Anopheles stephensi* larvae during March 2023 in Jigjiga, Ethiopia (visible as two orange squares with darker color in their center representing water). (B) Use of Google Earth Pro to digitize all visible construction pits (in this panel, a total of 5 pits are identified with a pin). (C) Distribution of the 101 construction pits visually identified in November 2022, 4 months prior to our sampling, within a rural-urban swath measuring 4.3 km^2^ and centered on the highway connecting Jigjiga with Somaliland (one of the busiest corridors in the region). (D) Kernel density estimate of the density of construction pits per hectare (color surface) and location of all identified pits (dots) using a bandwidth of 500m and a pixel size of 10m. Stars in C and D indicate the location of Jigjiga’s downtown.

### Political and economic development and *An. stephensi* invasion in Jigjiga

We consider that an unprecedented urban development boom in Jigjiga has been critical in favoring *An. stephensi* establishment and rapid spread. Jigjiga increased its built-up area from 4·2% in 1985 to 5·2% in 2005 to 24·0% in 2015, primarily driven by a change in status from zonal capital to regional capital, which opened political and economic opportunities, leading to high rural-urban immigration^26^. Since 2018, when the most transformative political reform of Ethiopia was enacted by the Ethiopian government, Jigjiga has seen an even larger population and urban footprint increase. The recent declaration by the national government of Ethiopia that 19·0% of commodity imports for the country ought to enter via port Berbera in Somaliland and transported through Jigjiga to the rest of the country^27^ led to an increased interest in investment and even higher immigration into the city^27^. Jigjiga’s population grew from 125,876 inhabitants in 2007 to more than 700,000 in 2020.

Since the 2018 political reform in Ethiopia, different groups began to accept Jigjiga’s new regional status as a safe regional hub, opening the window of opportunity to increased investment and business development^27^. Diaspora Somalis started to make investments, purchase land, and construct homes leading to a construction boom and increases in the price of land ^27^. New hotels, restaurants, as well as businesses are being built in preparation for the increased trade (and truck traffic) with port Barbera ^27^. As the city continues its unprecedented expansion, it is also increasingly facing critical water shortages (particularly during the dry season); the mean water accessibility of Jigjiga in 2016 was only 19·0% ^28^. In response to these water shortages, communities build cement cisterns to store water for domestic uses ^28^. Similarly, for building construction or brick manufacturing purposes, people in the town are accustomed to construct temporary construction pits lined with plastic sheet (Fig 1). During the dry season, water for construction pits is generally purchased and delivered in truck cisterns, which source the water from underground wells located outside the city. We can see evidence of the unprecedented construction boom in Jigjiga using historical satellite imagery (Fig 3). From the images one can see the dramatic expansion of construction pits in 2018 as well as the construction further along the periphery of the city. The sector went from 62-84 pits between 2016-2018 to 232 in 2020 and 192 in 2021, showcasing the rapid urban expansion of Jigjiga during that time (Fig 3). Construction pits are not only common practice in Jigjiga. In India, it is widely recognized that many *Anopheles stephensi* breeding sites are built into the finished structures of offices, homes and factories in urban areas ^29^. Less widely recognized, but also important as *An. stephensi* breeding sites, are the transient structures created during and as part of the construction process ^30^. Interestingly, this association between urban development and *An. stephensi* resembles the finding of cutaneous leishmaniasis outbreaks in association with urban growth and, specifically, construction sites in Israel settlements ^31,32^.

**Figure 3.**
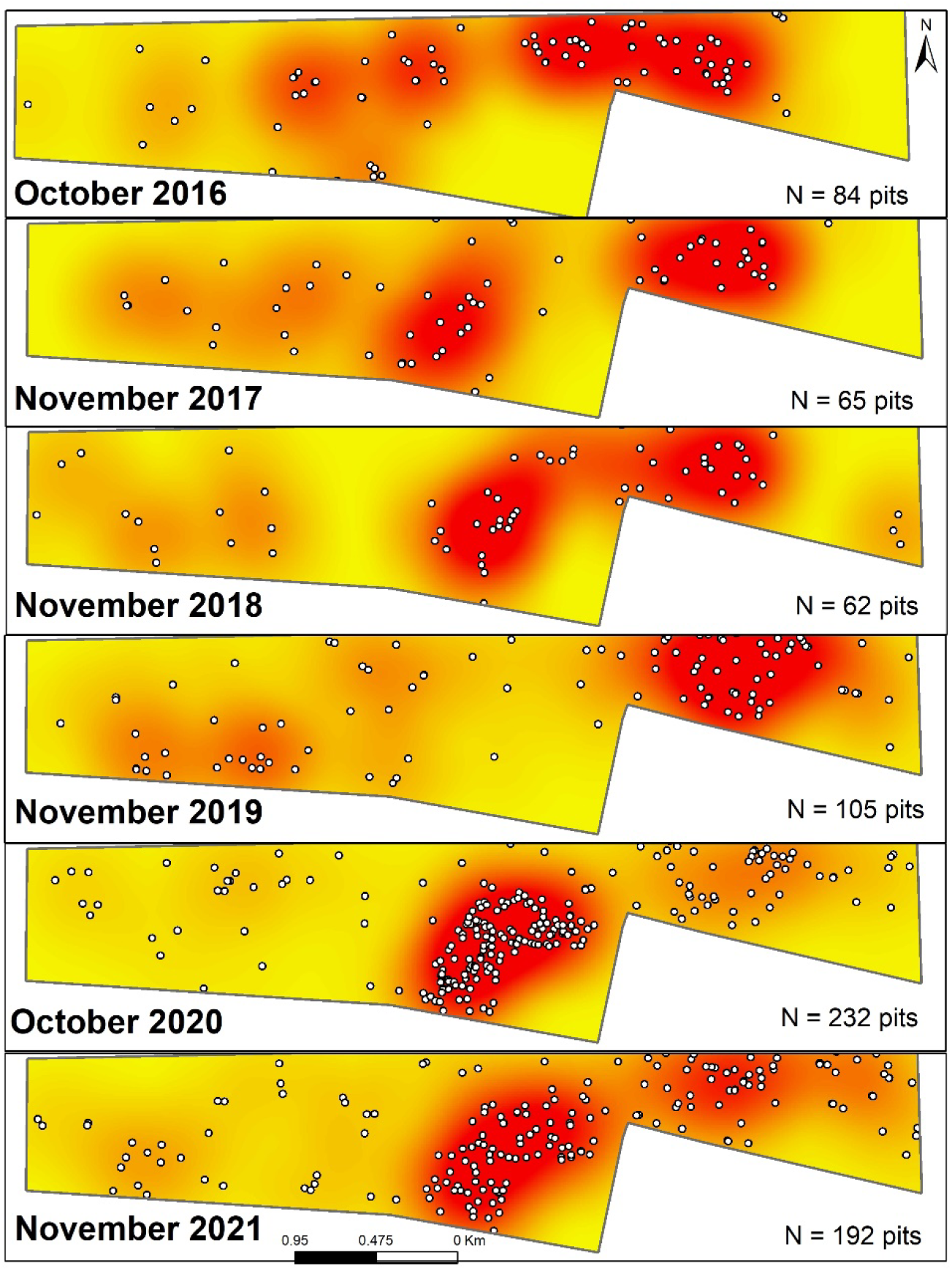
Historical sequence of the distribution of construction pits (white dots) in an urban-rural swath of Jigjiga, Somali region, Ethiopia. The dots represent digitized construction pits, observed through high-resolution satellite imagery historically archive in Google Earth. For each year we used October-November, as they were the months that had most complete information. The total number of pits per year on the area is listed on each panel.

### Opportunities for *An. stephensi* containment: larval source management of construction pits?

The finding of discrete and easily identifiable *An. stephensi* larval habitats in Jigjiga may provide a unique opportunity for immediate larval source management (LSM) and targeted control during the dry season, particularly with larviciding or biological control. A similar concept of ‘dry season LSM’ has been proposed for *An. gambiae* in semi-arid Kenya as an approach to maximize the effectiveness of larval control ^33^. An extensive list of larvicides prequalified by WHO for vector control exists ^34^. While temephos and Bacillus thuringiensis have shown important larviciding effect on *An. stephensi* from Ethiopia ^35^, they require frequent reapplication, which may be challenging given the number of construction pits that need to be treated. Long-lasting larvicide formulations, that could be potential candidates for control in large water volumes are Spinosad 7·48% DT (Clarke Mosquito Control Products, Inc.) and SumiLarv 2 MR (Sumitomo Chemical Co. Ltd., Japan). Spinosad DT is a tablet for direct application used at the dosage of 0· 5 mg/L AI (1 tablet/200 L) for control of container-breeding mosquitoes with a minimal expected duration of optimum efficacy of 4-6 weeks under field conditions ^36^. SumiLarv 2 MR is a 2 g plastic disc containing 2% (20 g AI/kg ± 25% w/w) pyriproxyfen used at the dosage of one disk in a water container with a volume of 40 L^37^. Long-lasting methoprene briquettes are commonly used for *Culex pipiens* control in catch basins in the US and, if prequalified, would provide an additional long-lasting tool since there is a 6-month extended release formulation (Altosid® 150-Day Briquets, Zoecon)^38^.

Given the water source and use, long duration of construction pits, and constant availability of water, a biological control option that can be considered is the use of larvivorous fish ^39^. Fish that feed on mosquito larvae have been widely used around the world in attempts to control malaria, other mosquito-borne diseases and mosquito nuisance biting ^39^ and could be used in this case as a “textbook” example ^40^. Locally native larvivorous fish exist near Jigjiga^39^. Furthermore, LSM in Jigjiga could include both larviciding and larvivorous fish if larvicides with low toxicity (Spinosad, metoprene or pyriproxyfen) are chosen. More importantly, our finding of high co-habitation between *An. stephensi* with *Culiseta spp*. and *Aedes spp*. mosquitoes provides a unique opportunity for integrated LSM across vectors, which can lead to important co-benefits and a higher justification for the implementation of such programs within Jigjiga and othe cities. Although for malaria typically the emphasis is on the protection of people inside their home (through deployment of ITNs and IRS). In the case of *An. stephensi* in Jigjiga, vectors could be controlled outside the house by conducting LSM during the dry season (the period when mosquito populations are lower and primary larval sites are easier to identify) to reduce the risk of vector establishment and further transmission malaria.

## Concluding remarks

The spread of *An. stephensi* in Africa may be facilitated in some Ethiopian cities by high urban immigration and an unprecedented construction boom, which is generating novel larval habitats that the vector exploits during the dry season to survive harsh environmental conditions. Our viewpoint emphasizes that the spread and persistence of *An. stephensi* in Jigjiga and other cities in Ethiopia is a planetary health problem that requires a holistic consideration of the environmental, social, and political changes that may be favoring the establishment and onward spread of this major threat to the elimination of malaria from sub-Saharan Africa.

## Acknowledgements

We thank the Jigjiga residents for allowing us to conduct this research. Also, we thank Jennifer Snyder, Bella Roeske and Cameron Goetgeluck for helping digitize the pit locations. EGHI Rapid Response Grant funding was provided by the Emory Global Health Institute to conduct this research. The content is solely the responsibility of the authors and does not necessarily represent the official views of the Emory Global Health Institute or the official policy or position of the Centers for Disease Control and Prevention.

